# Compartment-Resolved Matrisome Shifts During Postnatal Tendon Maturation in Mice

**DOI:** 10.64898/2026.11.03.764030

**Authors:** Benjamin Peterson, David Ceja Galindo, Maxwell McCabe, Jenna Galloway

## Abstract

Tendons acquire their load-bearing capacity during the first month of postnatal growth, driven by the extensive remodeling of the extracellular matrix (ECM). While the role of specific individual components to this process have been well characterized, a comprehensive assessment of how the tendon matrisome changes during this critical window is lacking. This, in part, is due to the challenges associated with solubilization of mature ECM, which is underrepresented by conventical proteomic approaches. To address this, we performed compartment-resolved proteomics on murine Achilles and plantaris tendons at postnatal days 0 (P0), P14, and P28. Sequential extraction methods leveraged a CHAPS-soluble fraction, capturing cellular and soluble matrisome proteins, from the insoluble mature ECM fraction, solubilized by guanidine hydrochloride and hydroxylamine hydrochloride and separately analyzed by label-free quantitative LC-MS/MS. Analysis shows that the two compartments followed distinct trajectories during maturation. The insoluble matrix was dominated by collagenous proteins, which increased from 95% to over 99% of the identified signal intensity, and its composition was largely established by P14. Within the soluble matrisome compartment, collagens increased from 50% to over 60% by P14, accompanied by dynamic regulation of proteins implicated in collagen processing (P4HA1, PCOLCE), crosslinking (LOX, LOXL2), and regulators of collagen fibril assembly and growth (DCN, BGN, FMOD, THBS3, THBS4). Beyond these established regulators, we observed dynamic regulation of less-characterized fibril-associated proteoglycans (KERA, OGN, CHAD), matricellular (COMP, POSTN, MATN2), and angiogenic regulatorily proteins (SERPINF1, ECM1). Together, this work establishes a comprehensive protein-level reference for postnatal tendon maturating, provides new insights into the molecular regulation of functional tendon development.

## 1. Introduction

Tendons are a dense connective tissue that transmits force between muscle and bone. The unique biomechanical capabilities of tendon are attributed to the highly ordered extracellular matrix (ECM), predominantly composed of type I collagen fibrils organized in a multiscale hierarchical arrangement^1^. In mice, these structure-function relationships are established in the first month of postnatal growth, driven by dynamic changes in the composition and organization of the collagenous matrix. During this period, tendons transition from a hyper-to hypocellular tissue as collagen fibrils are continuously deposited, crosslinked, and undergo longitudinal and lateral fusion to increase their length and diameter^23,4^. These changes coincide with the onset of weight-bearing activities and result in a rapid increase in tissue stiffness between postnatal days 14 (P14) and 21^4,5^. While the timing may vary, this transition is conserved across species^6,7^ and represents a critical stage in which the tendon acquires its functional capabilities.

In adult tendon, collagens account for 60-85% of the tissue dry mass, approximately 90% of which is type I collagen^8^. The remaining matrisome is composed by a mix of minor collagens, proteoglycans, glycoproteins, and matrix-modifying enzymes. Despite their low abundance, minority collages^9–12^ (e.g., Types III, V, XI, XII, and XIV) and small leucine-rich proteoglycans^13–15^ (e.g. decorin, biglycan, and fibromodulin) and have important roles in regulating collagen fibril nucleation, growth, and organization. During development, the timing of expression of these minority matrisome components are precisely coordinated and to modulate fibrillogenesis and matrix maturation. Yet, while the roles of individual components have been well characterized, a comprehensive assessment of how the matrisome changes during this critical period of maturation is still lacking.

Recent transcriptomic and epigenetic data has provided valuable insight into dynamic gene regulatory programs during these key stages of tendon development^16^. However, many structural matrix proteins are long-lived and heavily post-translationally modified, preventing their accumulated abundance from being fully captured by transcriptional profiling^17^. In contrast, proteomic approaches offer the ability to directly measure relative protein abundance across a variety of modification states, offering a more direct readout of matrix composition^18^. Despite this, proteomic studies of tendon have largely focused on adult tissue^19–22^, and the limited studies in developing tendon focused on a single developmental stage^23^ or isolated tissue compartments in non-murine models^24^.

A reason for this gap is likely due to the technical challenges in characterizing ECM-rich tissues. Mature fibrillar ECM (i.e., collagens and elastin) is intrinsically resistant to solubilization using conventional detergent-based extraction methods, often resulting in the underrepresentation of structural matrix proteins in standard proteomic datasets^18,25^. This challenge is further compounded during development, as post-translational modifications, such as procollagen processing or fibril crosslinking, dynamically alter protein solubility as the matrix matures. As a result, protein abundances quantified from a single-fraction extraction method may reflect changes in extraction rather than true changes in matrix composition and jeopardize comparisons across developmental stages.

To address these challenges, we performed compartment-resolved proteomics on isolated murine Achilles and plantaris tendons at P0, P14, and P28. Sequential extraction first isolated a CHAPS-soluble fraction, containing cellular and soluble matrisome proteins, from the insoluble mature ECM. This insoluble matrix was then solubilized by chemical digestion with guanidine hydrochloride and hydroxylamine hydrochloride, creating two distinct protein fractions representing distinct tissue compartments. Each fraction was then separately analyzed by label-free quantitative LC-MS/MS to enable a comprehensive assessment of the tendon matrisome across this critical window of tendon maturation.

## 2. Methods

### Animals

This study was conducted in accordance with protocols approved by the Intuitional Animal Care and Use Committee (IACUC) of Massachusetts General Hospital (Protocol #201300062). Adult C57BL/6J mice (Jax Strain #000664) were obtained from Jackson Laboratory and allowed one week of acclamation before breeding pairs were established. Females were checked for plugs to confirm conception and after 20 days breeding cages were checked twice daily (morning/afternoon) for pups to establish an accurate P0 timepoint. Mice were euthanized at postnatal (P) days 0 (P0), 14 (P14), or 28 (P28), for tissue proteomics or immunostaining following the described methods.

### Proteomics

#### Tissue Collection

At the designated timepoint, pups were euthanized according to AVMA guidelines of euthanasia based on stage. The skin was removed and the Achillies and plantaris tendons were micro-dissected from both hindlimbs. Tendons were cut above the calcaneal enthesis and below the myotendinous junction (**Fig. 1C**) to isolate the tendon midbody. Once isolated, the tendon was lightly scrapped with a micro-scalpel to minimize off-target tissue contaminates. Dissections were conducted in ice-cold 1x PBS with protease inhibitors (cOmplete^™^ EDTA-free, Roche). Following collection, samples were spun down, snap frozen in liquid nitrogen, and then stored at -80°C until proteomic characterization.

**Figure 1.**
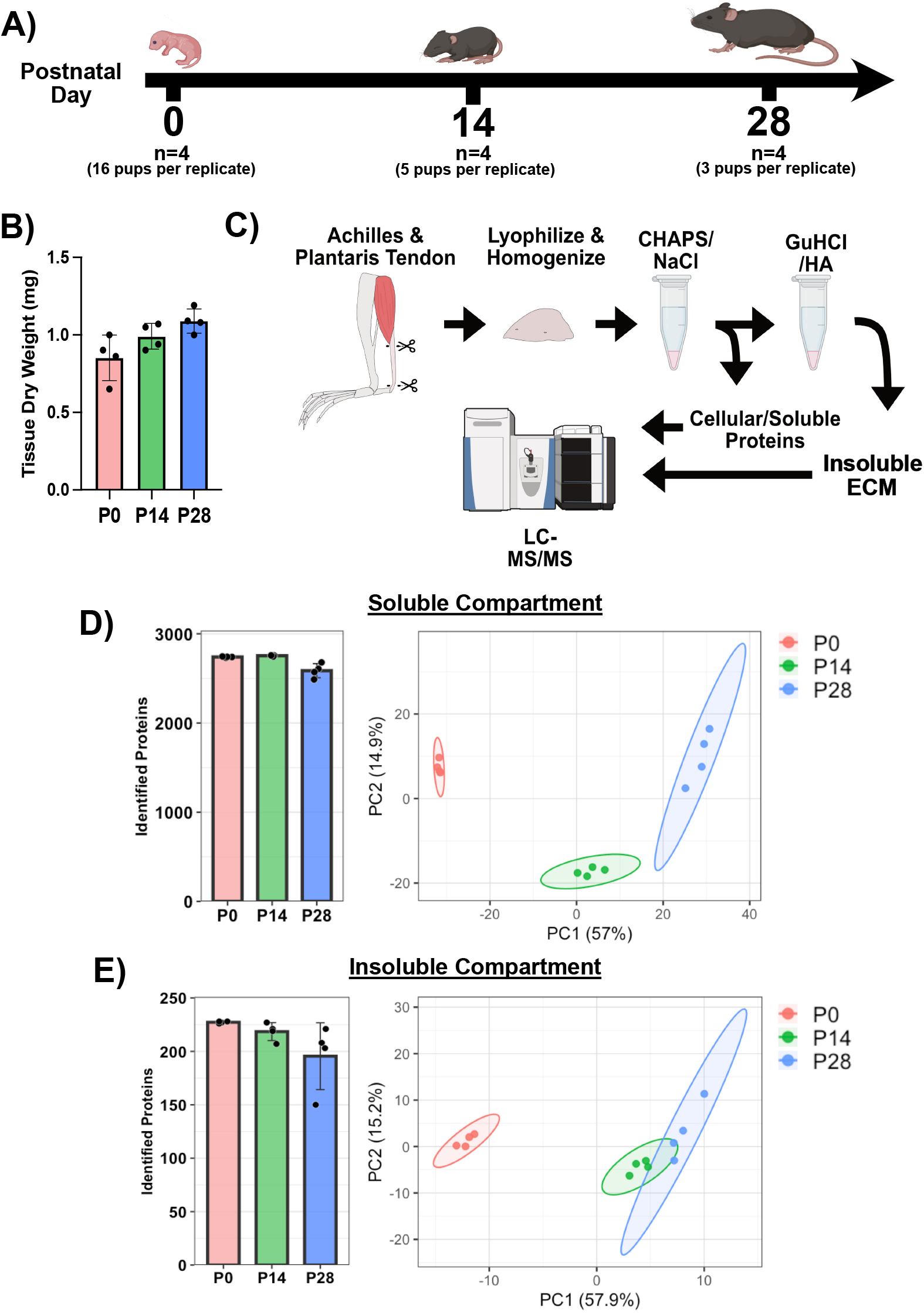
Overview of proteomic workflow. **A)** Timeline describing tissue collection from the murine hindlimb and multi-animal pooling strategy. **B)** The average tendon dry weight collected per timepoint. **C)** Flowchart describing the compartment-resolved proteomics procedure. **D & E)** The total identified proteins within the respective soluble/cellular and insoluble/extracellular compartments following label-free quantification (LFQ). Principle component Analysis (PCA) of the cellular and extracellular compartments. Of note, the cellular fraction showed distinctive cluster separation across all three timepoints along PC1. However, the extracellular fraction only showed clear separation between the early (P0) and late (P14 and P28) clusters.

#### Tissue Pooling

For each postnatal timepoint, four biologically equivalent replicates (n = 4) were collected for proteomic analysis (**Fig. 1A**). To ensure adequate quantities of protein were collected for LC-MS/MS, tendon tissue was collected from both hindlimbs and pooled across multiple pups *and* litters. To minimize litter-effects, pups from unique litters were evenly split across our four biologically equivalent replicates. For the P0 timepoint, tissue from 16 pups were collected per replicate across eight litters. For the P14 and P28 timepoints, a total of five and three pups were collected per replicate across four and three litters, respectively. Efforts were made to split sexes (i.e., males/females) evenly across replicates to mitigate sex-specific contributions and match the total amount of tissue dry weight (∼1mg) collected to minimize variability in tissue preparation.

#### Sample Preparation for Proteomic Analysis

Flash frozen tendon samples (pooled as described above) were lyophilized overnight using a FreeZone 4.5 L benchtop freeze dryer (Labconco #7750020) and milled to a fine powder prior to extraction. One milligram of material from each sample was combined with 100 mg of 1 mm glass beads (Next Advance #GB10) and homogenized in 200 μL/mg of decellularization buffer (50 mM Tris-HCl (pH 7.4), 0.25% CHAPS, 25 mM EDTA, 3 M NaCl, 1X Halt™ Protease Inhibitor Cocktail (Thermo Scientific #78438)) at power 8 for 3 min (Bullet Blender, Model BBX24, Next Advance, Inc.). Homogenate was then vortexed (power 8) for 20 min at 4°C, spun at 18,000 x g (4°C) for 15 min, and the supernatant was collected. Decellularization was repeated for a total of 3 washes, homogenizing and vortexing samples before each collection, and all washes were pooled to generate the cellular fraction. Pellets were then treated with freshly prepared hydroxylamine (HA) buffer (1 M NH_2_OH−HCl, 4.5 M Gnd−HCl, 0.2 M K_2_CO_3_, pH adjusted to 9.0 with NaOH) at 200 μL/mg of the starting tissue dry weight. Samples were homogenized at power 8 for 1 min and incubated at 45°C with shaking (1000 rpm) for 4 hours. Following incubation, the samples were spun for 15 min at 18,000 x g and the supernatant was removed before being stored at −80°C as the extracellular matrix (ECM) fraction until further proteolytic digestion with trypsin. All fractions were subsequently subjected to overnight enzymatic digestion with trypsin (1:100 enzyme:protein ratio) using a filter aided sample preparation (FASP) approach^26^ and desalted during Evotip loading (described below).

#### LC-MS/MS Analysis

Digested peptides (200 ng) were loaded onto individual Evotips following the manufacturer’s protocol and separated on an Evosep One chromatography system (Evosep, Odense, Denmark) using a Pepsep column, (150 µm inter diameter, 15 cm) packed with ReproSil C18 1.9 µm, 120Å resin. Samples were analyzed using the instrument default “30 samples per day” LC gradient. The system was coupled to the timsTOF Pro mass spectrometer (Bruker Daltonics, Bremen, Germany) via the nano-electrospray ion source (Captive Spray, Bruker Daltonics). The mass spectrometer was operated in PASEF mode. The ramp time was set to 100ms and 10 PASEF MS/MS scans per topN acquisition cycle were acquired. MS and MS/MS spectra were recorded from m/z 100 to 1700. The ion mobility was scanned from 0.7 to 1.50Vs/cm^2^. Precursors for data-dependent acquisition were isolated within±1Th and fragmented with an ion mobility-dependent collision energy, which was linearly increased from 20 to 59eV in positive mode. Low-abundance precursor ions with an intensity above a threshold of 500 counts but below a target value of 20000 counts were repeatedly scheduled and otherwise dynamically excluded for 0.4min.

#### Global proteomic data analysis

Data was searched using MSFragger v4.1 via FragPipe v22.0^27^. Precursor tolerance was set to ±12 ppm and fragment tolerance was set to ±25 ppm. Data was searched against UniProt restricted to *Mus musculus* reviewed sequences with added common contaminant sequences^28^ (17,340 total sequences). Enzyme cleavage was set to semi-specific trypsin for all samples. Fixed modifications were set as carbamidomethyl (C). Variable modifications were set as oxidation (M), oxidation (P) (hydroxyproline), deamidation (NQ), Gln->pyro-Glu (N-term Q), and acetyl (Peptide N-term). Label free quantification was performed using IonQuant v1.10.27 with match-between-runs enabled and default parameters. Cell and ECM fractions were searched separately for fraction-specific analysis. Results were filtered to 1% FDR at the peptide and protein level.

#### Data Processing and Differential Expression Analysis

For differential protein analysis, the individual soluble/cellular and extracellular protein fractions were analyzed separately to capture dynamic changes within their respective protein compartments during critical stages of tissue maturation. For each compartment, data sets were filtered such that proteins were retained only if they had non-zero LFQ intensity values in at least two out of the four biological replicates at every timepoint (P0, P14, and P28), with zeros converted to *NA* and no imputation applied. The filtered proteins were Log2 transformed, and each sample was median normalized (**Sup. Fig. 1**). Both the isolated protein fractions were processed independently under identical parameters unless otherwise specified.

To assess gross differences across maturation, principal component analysis (PCA) was conducted using *prcomp* (v4.4.3, R Studio) on the normalized datasets. Note, *prcomp* cannot handle ‘*NA’* values so PCA analysis was restricted to proteins fully identified across all samples. Differentially expressed proteins (DEPs) were assessed using *limma (v3*.*62*.*2, R Studio)*, fitting a linear model across developmental time periods with P0 as the reference. DEPs changing during postnatal maturation were identified via the moderated F-statistic and an adjusted p-value (adj. p) < 0.05, calculated by the Benjamani-Hochberg method. Pairwise comparisons (P14 vs P0, P28 vs P0, and P28 vs. P14) were calculated via the same linear fit and proteins with an adj. p < 0.05 and |log_2_-fold Change| > 0.58 (|FC| > 1.5) were considered differentially significant. Downstream analysis focused on matrisome-associated proteins, cross-referencing identified proteins with annotations from the MatrisomeDB. Gene set enrichment analysis (GSEA) was conducted using *clusterProfiler* (v4.14.6) to identify enriched biological pathways between timepoints, ranking proteins by their moderated t-statistic. For all plots, the protein’s corresponding gene name was utilized.

## 3. Results

### 3.1 Compartment-resolved proteomics reveals distinct protein trajectories during maturation

To capture protein-level remodeling during postnatal tendon maturation, we performed compartment-resolved proteomics on isolated Achilles/Plantaris tendons from the murine hindlimb at P0, P14, and P28 (**Fig. 1A**). Sequential extraction methods separated the bulk tissue into distinct soluble/cellular and insoluble/extracellular matrix protein fractions^18^. After processing, 2802 and 228 unique proteins were identified within the soluble/cellular and extracellular compartments, respectively. Principle component analysis (PCA) revealed unique clustering behaviors across compartments as a function of developmental time. Specifically, the soluble/cellular fraction reports separation of the P0, P14, and P28 clusters along the PC1 axis (**Fig. 1D**), indicating a shifting protein composition across the full postnatal developmental window. Conversely, the ECM fraction only shows separation between the early P0 cluster from later timepoints (P14 and P28), which were grouped more closely together. Of note, we observed tighter in-group clustering behavior of samples across both protein fractions at earlier developmental stages, likely attributed to their increase sample pooling.

### 3.2 The soluble matrisome undergoes extensive and sustained remodeling during postnatal maturation

Differential expression analysis of the soluble/cellular protein fraction identified 1,808 unique DEPs across postnatal maturation (adj. p < 0.05 **Fig. 2A**). More stringent individual pairwise contrasts (adj. p < 0.05, |Log_2_FC| > 0.58), report 1,139 DEPs between P0 vs P14, 1,626 at P0 vs P28, and 991 at P14 vs. P28 (**Fig. 2B**). Notably, persistent quantities of DEPs were observed between the P28 and P14, suggesting dynamic protein expression across the full postnatal window and consistent with the principal component analysis (**Fig. 1D**). Gene set enrichment analysis (GSEA) of the soluble/cellular proteome reports distinctive shifts within biological pathways, including the activation of pathways associated with connective tissue, skeletal tissue, and cartilage development (**Supp. Fig 2**). We further observed a suppression in pathways associated with DNA replication and cell cycling, indicative of a decrease in proliferative behavior with maturation.

**Figure 2:**
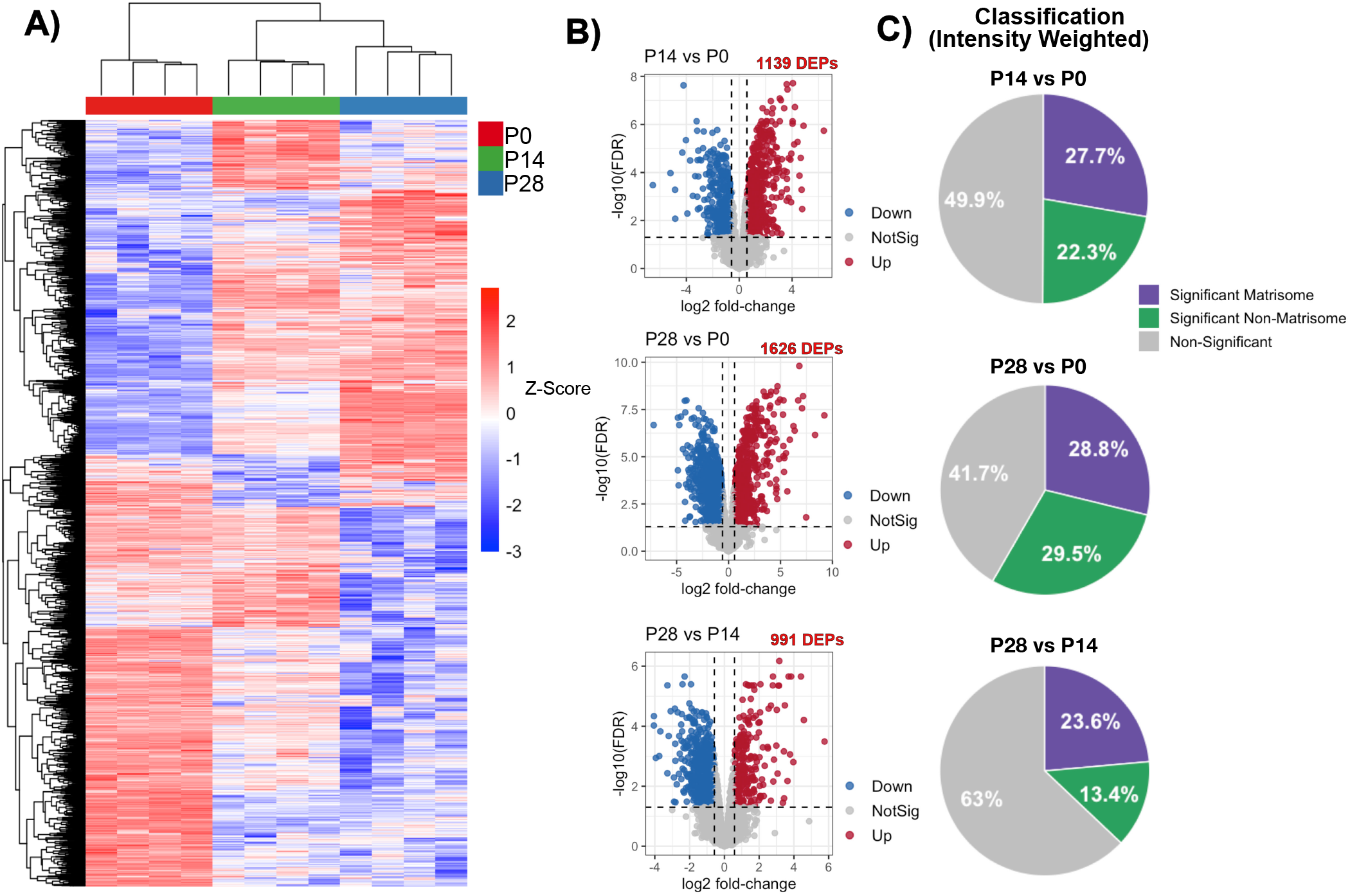
Differential expression analysis of the soluble/cellular fraction. **A)** Clustered heatmap of the 1808 identified differentially expressed proteins (DEPs) (adj. p < 0.05). **B)** Volcano plots of signficant DEPs between P14 vs P0 (1139 DEPs), P28 vs P0 (1626 DEPs), and P28 vs. P14 (991 DEPs). Significance set at adj. p < 0.05 and |log2-Fold Change| > 0.58. **C)** Identified proteins classified according to their contributions to the tissue matrisome within each contrast, weighted by LFQ intensity. Classifications assigned utilizing annotations within MatrisomeDB.

For the purposes of this study, we focused on specific proteins within the soluble/cellular proteome associated with the establishment of the extracellular matrix. Utilizing annotations within MatrisomeDB, all identified proteins were classified according to their association to the matrisome, significance (adj. p < 0.05, |Log_2_FC| > 0.58), and represented according to their LFQ-weighted contribution to the total proteome for each contrast. While the number of significant proteins varied between contrasts, the weighted contribution of matrisome DEPs was persistent across maturation, representing approximately a quarter of the total proteome **(Fig. 2C)**.

Identified proteins were categorized into subclassifications based on their structural role or regulatory functions within the matrisome (**Fig. 3A**). At P0, collagens accounted for 50% of matrisome protein signal in the soluble fraction, with the remainder distributed across ECM regulators (13.48%), ECM-affiliated proteins (12.54%), glycoproteins (12.07%), proteoglycans (11.4%) and secreted factors (0.56%). By P14, the fraction of matrisome signal assigned to collagen increased to 63.2% and remained comparable at P28 (62.6%). With maturation, we observed a progressive decline in the compositional fraction of identified ECM regulators, ECM-affiliated, and glycoprotein categories, and an increase in the resident proteoglycans (**Fig. 3A**).

**Figure 3.**
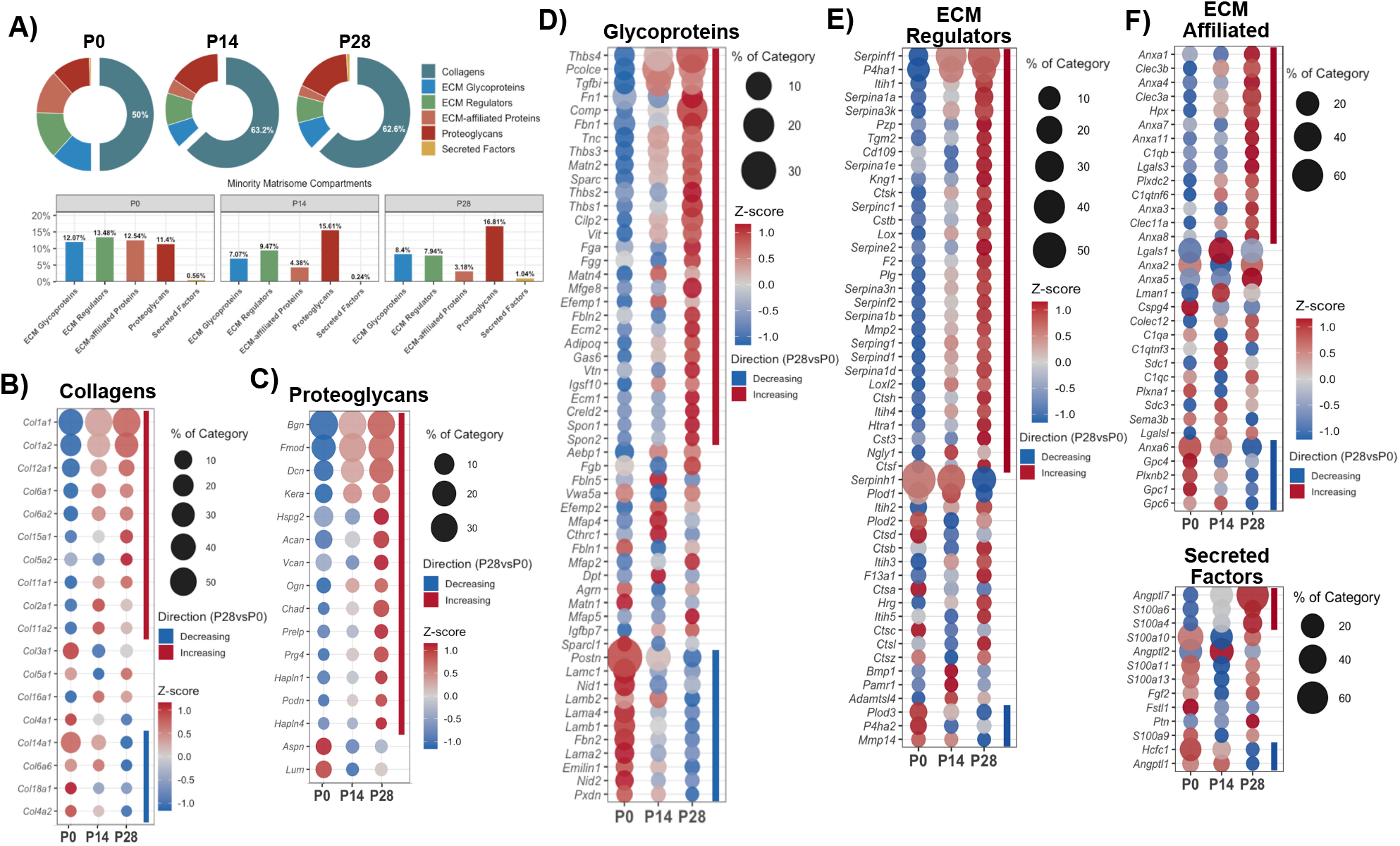
Remodeling of the soluble/cellular tendon matrisome during postnatal maturation. **A)** Matrisome composition with age, shown as the intensity-weighted proportion of each matrisome sub-category. Dot plots of the category-resolved protein dynamics for all identified **B)** Collagens, **C)** Proteoglycans, **E)** ECM Regulators, **F)** ECM-Affiliated Proteins and Secreted Factors. Color encodes the row z-score of normalized abundance across development. Dot size represents the components percentage contribution to the total protein abundance within each category at each age. The colored bar indicates whether a protein significantly increased or decreased as a function of maturation (P28 vs. P0). Significance was defined as *adj. p* < 0.05 and |Log_2_ fold change| > 0.58. Protein matrisome classifications were assigned using MatrisomeDB.

### 3.3 Dynamic regulation of collagen synthesis, processing, and assembly

Given that matrix-associated proteins account for a prominent share of the differentially expressed proteome, we next aimed to investigate their dynamic relationships during tendon maturation. Critically, the soluble protein fraction extracted using CHAPS, a non-denaturing zwitterionic detergent, is incapable of solubilizing mature, crosslinked collagens^29^. As a result, the collagenous populations measured here (**Fig. 3B**) best reflect dynamic changes in immature procollagen and serve as a proxy for active collagen synthesis from resident tenocytes. We observed a net increase in the collagenous fraction (**Fig. 3A**) at P14 and P28 in comparison to P0, consistent with an increase in collagen synthesis at later maturation stages. This shift was driven predominantly by an increase in collagen Type I (COL1A1, COL1A2), though other fibrillar (COL5A2, COL11A2, and COL2A1), non-fibrillar (COL6A1, COL6A2), FACIT (COL12A1), and multiplexin (COL15A1) collagen subtypes also increased with age (**Fig. 3B**). A separate subset, including COL14A1, COL6A6, COL18A1, and COL4A2 were comparatively enriched at P0 and decreased with maturation. Finally, several collagens (COL3A1, COL5A1, COL16A1, and COL4A1) were detected at similar levels across all developmental stages but were not significantly altered.

Matching the rise in fibrillar collagens, we observed coordinated regulation of collagen processing components (**Fig. 3E**). The prolyl 4-hydroxylase subunit alpha 1 (P4HA1), which facilitates the hydroxylation of proline in the collagen monomer^30^, increased with maturation in conjunction with the rise in the collagenous fraction. Interestingly, we found P4ha2 was inversely correlated, being enriched at P0 and decreasing with maturation, indicating an isoform-specific shift in the hydroxylation machinery during development. Lysyl hydroxylases (PLOD1, PLOD2, and PLOD3) were also identified, which facilitate the hydroxylation of lysine residues required for enzymatic crosslink formation^31,32^. PLOD1 and PLOD2 were actively expressed across all timepoints but were not significantly altered, whereas PLOD3 was enriched at P0 and decreased with maturation (**Fig. 3C**). This shift occurred in conjunction with a rise in procollagen C-proteinase enhancer (PCOLCE) over time (**Fig. 3D**), which promotes proteomic cleavage of pro-collagen C-terminus. Finally, we observed an increase in lysyl oxidase components LOX and LOXL2 with maturation (**Fig. 3E**), which interact with lysine and hydroxylysine residues of collagen and elastin monomers to initiate crosslinking^32^.

Proteins associated with downstream fibril assembly and organization were also altered with maturation. Among the detected glycoproteins, we observed an increase in the thrombospondins THBS3, THBS4, and COMP (THBS5) (Fig.3D), a family of oligomeric matricellular proteins capable of binding to collagen and mediate fibril assembly^33–36^. Similarly, the matricellular proteins SPARC, which has been implicated in the regulation of procollagen processing and mediating cell-matrix interactions^37^, increased during development. Small-leucine-rich proteoglycans (SLRPs) Decorin (DCN), Biglycan (BGN), and Fibromodulin (FMOD) were also highly expressed, and increased with maturation (**Fig. 3C**). Interestingly, we further identified other SLRPs, such as Keratocan (KERA) and Osteoglycan (OGN), which additionally increased with age.

### 3.4 Supporting Matrisome components shift with maturing matrix composition

Beyond collagen synthesis and assembly, maturation was accompanied by changes in matrisome components associated with the basement membranes, vascular development, and cell-matrix adhesions. Laminin subunits, including α1 (LAMA1), α4 (LAMA4), β1(LAMB1), and γ1 (LAMC1), were found enriched at birth (P0) and decreased with age (**Fig. 3D**). Laminins operate as core structural components of the underlying basement membrane and contribute to cell signaling via focal adhesions^38^. Their decline with maturation paralleled the reduction of basement membrane associated collagens COL4A2 and COL18A1 (**Fig. 3B**). Notable, the laminin subunits α4, β1, and γ1 form the laminin-411 complex, which serves as a principal component of the vascular basement membrane^39^, suggesting reduced vascularization with ECM maturation. Consistent with this, we observed a significant increase in SERPINF1 (**Fig. 3E**), a known negative regulator of angiogenesis^40^. Conversely, extracellular matrix protein 1 and 2 (ECM1, ECM2) increased with maturation, which positively regulates the formation of new blood vessels^41^. Together, implicating both pro- and anti-angiogenetic behavior are regulated during this period of tendon growth and development.

Proteins involved in cell-matrix interactions and the underlying supporting matrix organization were also dynamically altered. Periostin (POSTN), a matricellular protein that mediates cell adhesions^42,43^, was the most enriched glycoprotein at P0 and decreased with maturation. This pattern was confirmed by immunostaining (**Sup. Fig. 3A**). Among the microfibrillar proteins that coordinate elastin assembly^44,45^, we found fibrillin-2 (Fbn2) was enriched at birth and decreased with maturation, while fibrillin-1 exhibited the inverse (**Fig. 3D**). We further identified an increase in numerous supporting glycoproteins which contribute to cell-matrix interactions, predominantly driven by an increase in the fractional composition of fibronectin (FN1) and tenascin-C (TNC) (**Fig. 3D**). Similarly, we observed an increase in MATN-2 and -4, extracellular matrix adaptor proteins that mediate interactions between collagen fibril and other matrix components^46^, such as decorin, and a variety of lesser expressed SLRPs.

### 3.5 The ECM undergoes front-loaded compositional remodeling during tendon maturation

Following extraction of the cellular/soluble protein fraction, the remaining tissue was processed with guanidine hydrochloride and hydroxylamine hydrochloride, to enable the digestion, solubilization, and characterization of the mature extracellular matrix. Analysis of the ECM fraction identified 228 unique differentially expressed proteins (DEPs) across the three developmental timepoints (adj. p < 0.05, **Fig. 4A**). Pairwise contrasts (**Fig. 4B**) showed that majority of the significantly different (adj. p < 0.05, |log2FC| > 0.58) proteins were relative to the initial P0 baselines, with 111 DEPs between P0 vs. P14 and 137 DEPs between P0 vs. P28. Conversely, only 44 DEPs were identified between P14 vs. P28, mirroring trends observed within the PCA (**Fig. 1E**). The resulting DEPs were then classified according to their significance, matrisome association, and weighted by LFQ intensity to reflect the proportion of the proteome that shifting with maturation (**Fig. 4C**).

**Figure 4:**
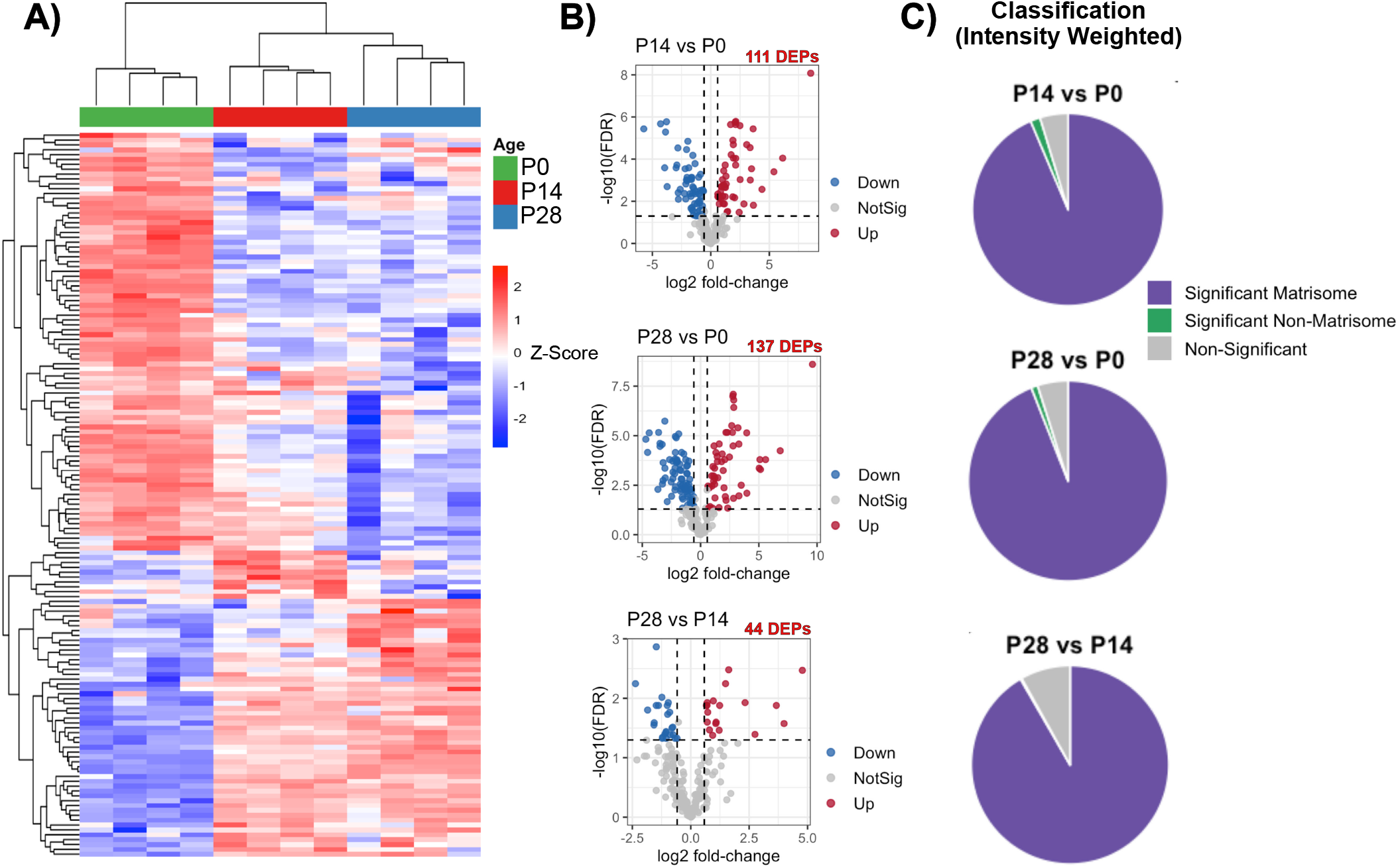
Differential expression analysis of the insoluble matrix fraction. **A)** Clustered heatmap of the 228 identified differentially expressed proteins (DEPs). **B)** Volcano plots of DEPs between identified P14 vs P0 (111 DEPs), P28 vs P0 (137 DEPs), and P28 vs. P14 (44 DEPs). Significance set at adj. p < 0.05 and log2-Fold Change > 0.58. **C)** Classification of proteins identified within each individual contrast, weighted by it’s LFQ intensity. Classification assigned according to annotations within MatrisomeDB and if differentially expressed between contrasts.

To investigate changes in protein composition, matrisome proteins were classified into distinct matrisome categories (**Fig. 5A**). As expected, the extracellular proteome fraction was dominated by collagenous components comprising 95% of identified matrisome protein signal at P0 to 98.9% and 99.2% at P14 and P28, respectively. This shift was driven by an increase in fibrillar collagens Type I (a1, a2), II (a1), and XI (a1, a2), along with multiplexin (COL18A1, COL15A1) and network-forming (COL4A2, COL10A1) collagens (**Fig. 5B**). Several fibrillar collagens (COL3A1 and COL5A1/A2) and fibril-associated collagen with interrupted triple helices (FACIT) (COL12A1, COL14A1, COL16A1) were also identified, but their relative composition unchanged with maturation.

**Figure 5.**
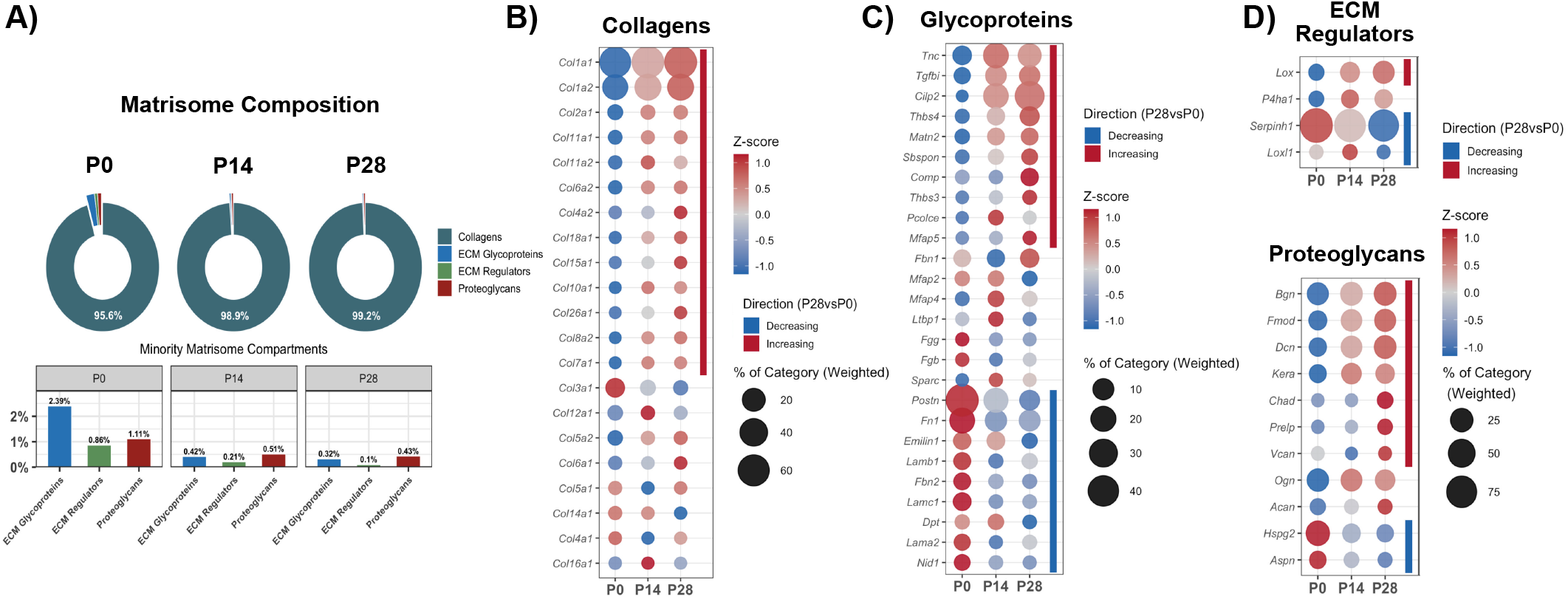
Remodeling of the Achilles/Plantaris insoluble fraction matrisome during postnatal maturation. **(A)** Insoluble matrisome composition at each age, show as the intensity-weighted proportion of each matrisome sub-category. Dot plots of the category-resolved protein dynamics for all identified **(B)** Collagens, **(C)** Glycoproteins, **(D)** ECM Regulators and Proteoglycans. Color encodes the row z-score of normalized abundance across development. Dot size represents the components percentage contribution to the total protein abundance within each category at each age. The colored bar indicates whether a protein significantly increased or decreased as a function of maturation (P28 vs. P0). Significance was defined as *adj. p* < 0.05 and |Log_2_ fold change| > 0.58. Protein matrisome classifications were assigned using MatrisomeDB.

Non-collagenous matrisome components were predominately identified and enriched within the soluble/cellular fraction (**Fig. 3**), but even at lower relative amounts we did detect significant populations within the insoluble ECM fraction (**Fig. 5A**). The limited detection and diversity of these minority compartments likely reflect residual or tightly collagen-associated components that persist through the cellular/soluble extraction.

## 4. Discussion

In this study, we used compartment-resolved proteomics to characterize protein-level remodeling of the murine Achilles/Plantaris tendons during the first month of postnatal maturation. This period of growth captures dynamic changes in tendon structure-function behavior and the acquisition of robust tensile properties^4^, critical for the onset of load-bearing actives (i.e., walking)^5^. While prior work has characterized individual matrix components during this period, a comprehensive assessment of the tendon proteome has been lacking. To address this, we used a sequential extraction method that leverages the differential solubility of the proteome to enrich targeted protein compartments, which were then analyzed separately to maximize detection sensitivity. Analysis of the two compartments showed distinct trajectories with maturation. The composition of the insoluble fraction, representing the mature ECM, was largely established by P14 (**Fig. 1E**), whereas the soluble proteome continued to change extensively through the full maturation period **(Fig. 1D)**. This suggests that the cellular programs that produce, organize, and remodel the matrix remain highly active well after the ECM relative composition is established.

Given our focus on matrix development, we centered our analysis on significant matrisome and matrisome-associated proteins, which accounted for 23.6 – 28.8% of intensity-weighted proteome changing with maturation in the soluble fraction. Critically, because CHAPS does not readily solubilize mature, crosslinked collagens^29^, the collagens detected within this fraction most likely reflect immature procollagen. Consistent with this, collagen components increased from 50% to over 60% of the soluble matrisome by P14, coinciding with an increase in procollagen processing proteins, particularly P4HA1 (**Fig. 3D-E**). *P4ha1* encodes the catalytic α1 subunit of collagen prolyl 4-hydroxylase, which hydroxylates proline residues within in Gly-X-Y sequence of various collage types and is required for procollagen assembly and secretion^30,31^. P4HA1 abundance is further associated with overall collagen biosynthesis capacity^31^, supporting the increase in collagen synthesis and accumulation observed at later postnatal stages (**Fig. 3B, Fig. 5B**). These finding align with previously published transcriptomic data, which found that *Col1a1* and *Col1a2* mRNA expression peaked at P14^16^. Beyond synthesis, collagen requires additional extracellular processing to facilitate downstream self-assembly. Notably, the primary procollagen C-proteinase BMP-1^472^ did not increase with maturation (**Fig. 3E**), despite the increase in upstream procollagen synthesis components. However, the BMP-1 enhancer PCOLCE expression was dynamically upregulated during this window (**Fig. 3X**), suggesting that C-terminal processing capacity may be regulated through enhancer availability rather that protease abundance, but warrants further investigation.

Additionally, machinery which coordinates the crosslinking of collagenous fibrils were identified. The lysyl hydroxylases PLOD1 and PLOD2, which facilitate the intracellular hydroxylation of lysine residues, remained highly abundant throughout maturation. PLOD2 is of particular interest, as it has been specifically implicated in hydroxylation of telopeptide lysines, a modification required for the formation of mature trivalent hydroxylysyl pyridinoline (HP) and lysyl pyridinoline (LP) crosslinks^47^. Follow secretion of the procollagen monomer, lysyl oxidases catalyze the lysine and hydroxylysine residues into reactive aldehyde species, initiating crosslink formation between adjacent collagen molecules^32^. Both LOX and LOXL2 were identified and increased with maturation (**Fig. 3E**), suggestive of an upregulation in crosslinking at later postnatal stages. These findings are consistent with previous work identifying the LOX family as an essential mediator of tendon biomechanical development^6^. However, it’s important to note that lysyl oxidases are secreted as an inactive proenzyme and our methods do not differentiate between the inactive and active form.

While type I collagen was the dominant collagen, we identified a diverse subset of minority collagens which have been implicated in regulating distinct stages of fibril assembly. Among these, the regulatory fibrillar collagens COL5A2, COL11A1, and COL11A2 increased with maturation, consistent with their roles in modulating fibril nucleation and diameter. In contrast, COL3A1 was detected at all timepoints but showed no changes. FACIT collagens showed divergent regulation, with COL14A1 decreasing and COL12A1 increasing with maturation. These patterns agree with genetic knockout studies demonstrating collagen XIV regulates the onset of fibril growth during earlier postnatal periods^12^, whereases collagen XII is required predominantly at later postnatal stages and contributes to the emergence of biomechanical capabilities^48^. Interestingly, we’ve also identified a number lesser-known collagen sub-types within both insoluble fraction that increased with maturation, including collagen types XVIII, XV, and XXVI (**Fig. 5B**). However, their contribution to tendon maturation, if any, are unclear.

Regulators of higher-order fibril assembly and growth were also found to be upregulated with maturation. The thrombospondins THBS3, THBS4, and COMP (THBS5) increased with maturation and are known to modulate fibril spacing and fusion behavior^33^. Interestingly, COMP has been implicated in the functional specialization of tendon, with higher expression in load-bearing tendons^49^. Mutations in COMP have also been linked to pseudoachondroplasia, a form of skeletal dysplasia characterized by ligament laxity and joint hypermobility^50^. The increased presence of COMP in the Achilles/Plantaris observed here may reflect physiological specification or adaptations of the tendon matrix in response to rising mechanical demand following the onset of load-bearing.

Consistent with prior literature, we identified an increased presence of the SLRPs Decorin (DCN), Biglycan (BGN), and Fibromodulin (FMOD), which regulate lateral and longitudinal fibril fusion during development^13,13,14^. More unexpectedly, we observed an increase in lesser known SLRPs during this maturation window including keratocan (KERA), osteoglycin (OGN) and chondroadherin (CHAD), which have been implicated in regulating fibrillogenesis but largely in other tissues, including cornea^51^, skin^52^, cartilage^53^, and bone^54^. While these SLRPs have been identified in mature tendon^20,24,55,56^, their contribution to tendon development is largely unknown. Similarly, we identified an increase in extracellular adaptor proteins MATN2 and MATN4, which bridge proteoglycans and collagen fibrils to form a filamentous network within the extracellular matrix^46^. While their role in tendon is still largely unclear, MATN2 has been implicated in regulating TGFB/Smad signaling^57^ and as an essential mediator of muscle regeneration^58^.

The soluble proteome fraction also captured broader changes in the tissue microenvironment. We observed a decline in laminin subunits with maturation, including subunits which specifically contribute to laminin-411, a primary component of the vascular basement membrane^39^. Concurrently, the angiogenesis inhibitor SERPINF1 (PDEF) was highly abundant and increased with maturation^40^. Although the progressive reduction in vascularity of maturing tendon is well recognized^59^, the molecular regulators of this transition are poorly characterized, and SERPINF1 is a compelling candidate for further investigation.

## 5. Conclusion

In summary, this study provides a compartment-resolved view into the postnatal tendon matrisome during critical stages of maturation in which tendon acquires its load-bearing capacity. Our data suggests that the deposited matrix is largely established by P14, but the cellular machinery that modulates collagen synthesis, processing, and remodeling remain highly dynamic throughout the first month. Beyond confirming known regulators of tendon development, our data has also identified less-characterized candidates that may contribute to tendon maturation and require further investigation. Further, these findings establish a protein-level baseline for tendon development that may inform strategies to improve the maturation of tissue engineering constructs or novel regenerative strategies.

### Limitations

Several limitations should be considered when interpreting these data. First, while valuable, sequential extraction does not achieve perfectly clean separation between compartments and risks carry-over between fractions. Further, solubility-based partitioning can bias detection of components that are poorly soluble or when solubility changes dynamically during development. For example, elastin, a major extracellular component of the Achilles tendon^60^, was filtered out during data processing due to a lack of detection at P28 and likely reflects a reduction in solubility at later maturation stages. Consistent with this, we observed an increase in components that support elastin assembly (FBN2) at later maturation stages. Second, label-free quantification is semi-quantitative and relationships in protein composition are relative to the total compartment. As a result, an apparent decrease in a given protein may reflect the accumulation of other proteins, such as the expanding collagen fraction, rather than a reduction in its absolute abundance or expression. Additionally, a high degree of animal pooling was required and makes it difficult to assess inter-animal variability. Third, although compartment-resolved extraction improved detection, some low-abundance regulatory proteins were still not detected. For example, while we readily identified procollagen C-proteinases (BMP1), we didn’t identify the complementary N-terminal proteases (ADAMTS). It’s unclear if this reflected protein degradation during prep or a limitation in achievable proteome depth, even with the multi-fraction employed here. Finally, our analysis was performed on bulk tissue and therefore cannot resolve cell-type specific contributions to the proteome. Similarly, we may have observed tissue contamination from the myotendinous junction, enthesis, or residual blood within the tissue at collection, despite our best efforts to consistently collect from the mid-body of the tendon. Additionally, this study only profiled three timepoints and finer temporal sampling may be required to better resolve preside contributions to changes within tissue maturation.

