## Supplementary material for "Compartment-Resolved Matrisome Shifts During Postnatal Tendon Maturation in Mice": Supplmental Figures

### 7. Supplemental Information

##### *Immunostaining*

Following euthanasia, the skin was removed from the murine hindlimbs and fixed overnight in 4% PFA at 4$^{oC}$ on an orbital shaker. Limbs were briefly washed in PBS prior to decalcification in 0.5M EDTA, pH 8.0. Decalcification was conducted on the P0, P14, and P28 hindlimbs for 1 day, 5 days, and 10 days, respectively. Limbs underwent paraffin infusion, embedded for sagittal sections, and sectioned at a thickness of 5μm. For immunostaining, sections were deparaffinized and underwent antigen retrieval in a Tris-EDTA buffer (Abcam, ab93684) overnight at 60$^{oC}$. Endogenous peroxidases were quenched using BLOXALL (Vector Labs) prior to blocking. Slides were incubated with the primary antibody (**Supp. Table 1**), diluted in blocking buffer, overnight at 4$^{oC}$. Sections subsequently incubated with the goat anti-rabbit HRP (Vector Labs, MP-7451) and detected using DAB substrate (Vector Labs, SK-4105). Slides were counterstained using Hematoxylin QS (Vector Labs, H-3404) before dehydration, clearing, and mounting.

**Table 1:** Antibodies

| **Primary Antibodies** | **Manufacturer** | **Dilution** |
|---|---|---|
| *Rabbit anti-COMP (ab231977)* | Abcam | 1:200 (IHC) |
| Rabbit anti-Periostin (ab215199) | Abcam | 1:200 (IHC) |
| ***Secondary Antibodies*** | **Manufacturer** | **Dilution** |
| ImmPress HRP Goat Anti-Rabbit | VectorLabs | |

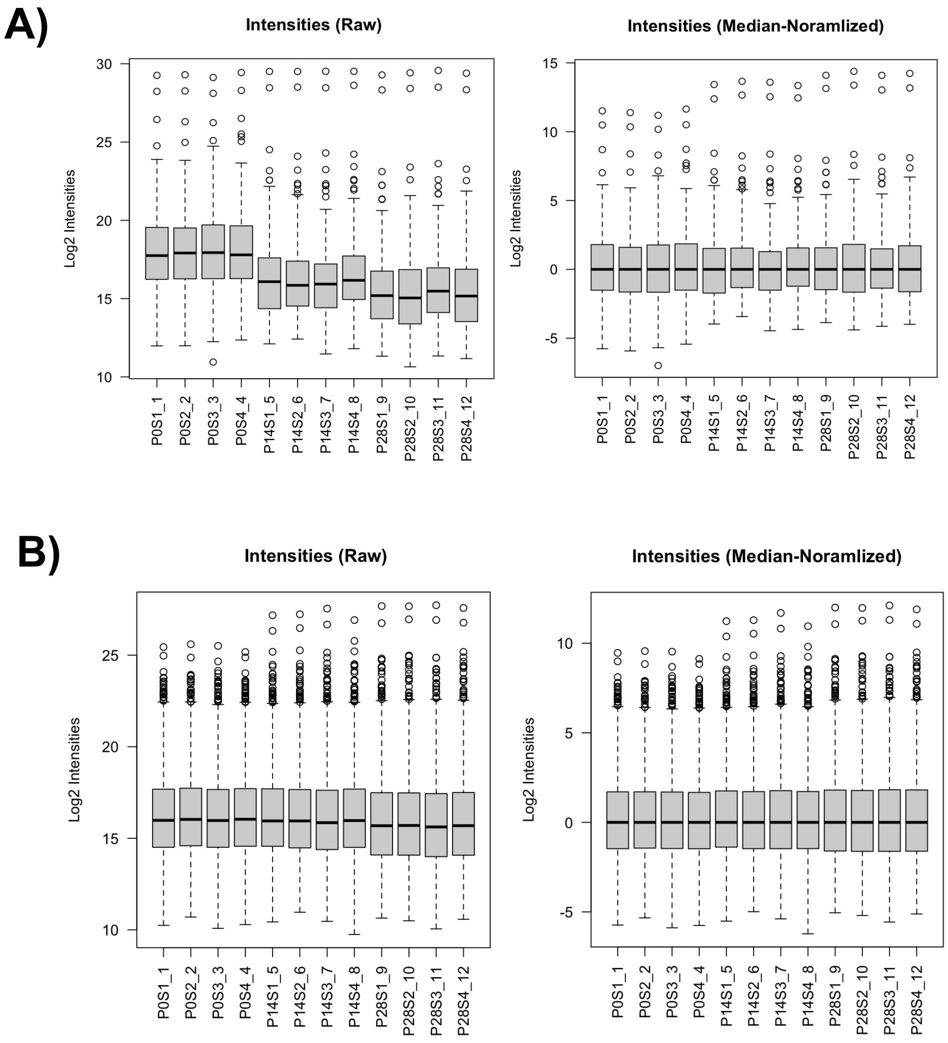

**Supplemental Figure 1**: Raw and median-normalized LFQ intensity values of the **A)** soluble/cellular and **B)** insoluble/extracellular matrix compartments.

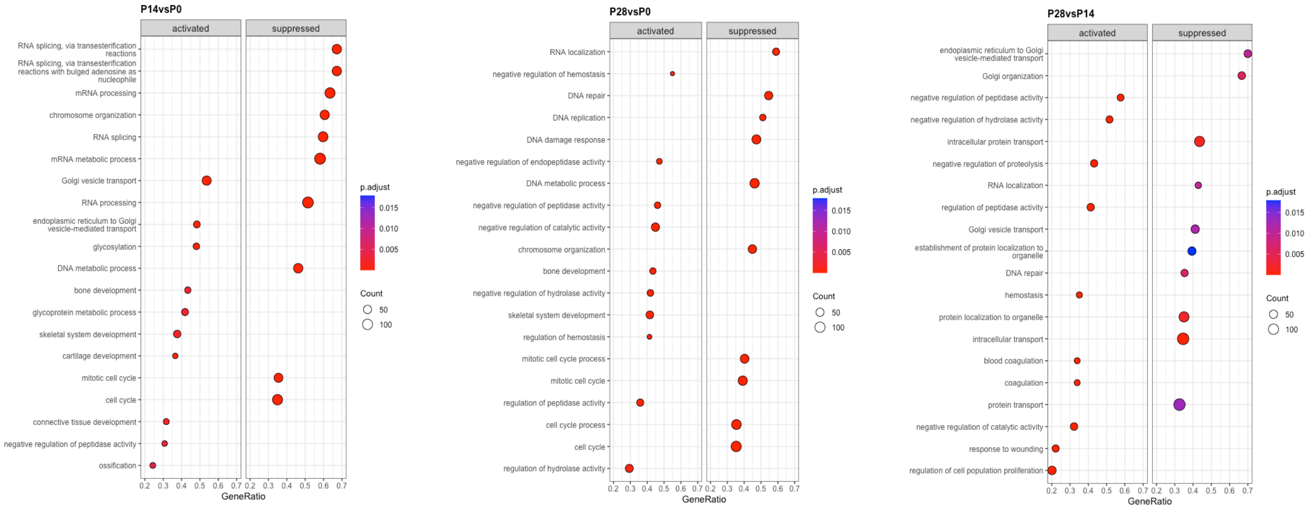

**Supplemental Figure 2**: Gene set enrichment analysis (GSEA) between developmental contrasts of the full soluble protein fraction. Proteins were ranked according to the moderated F-statistics.

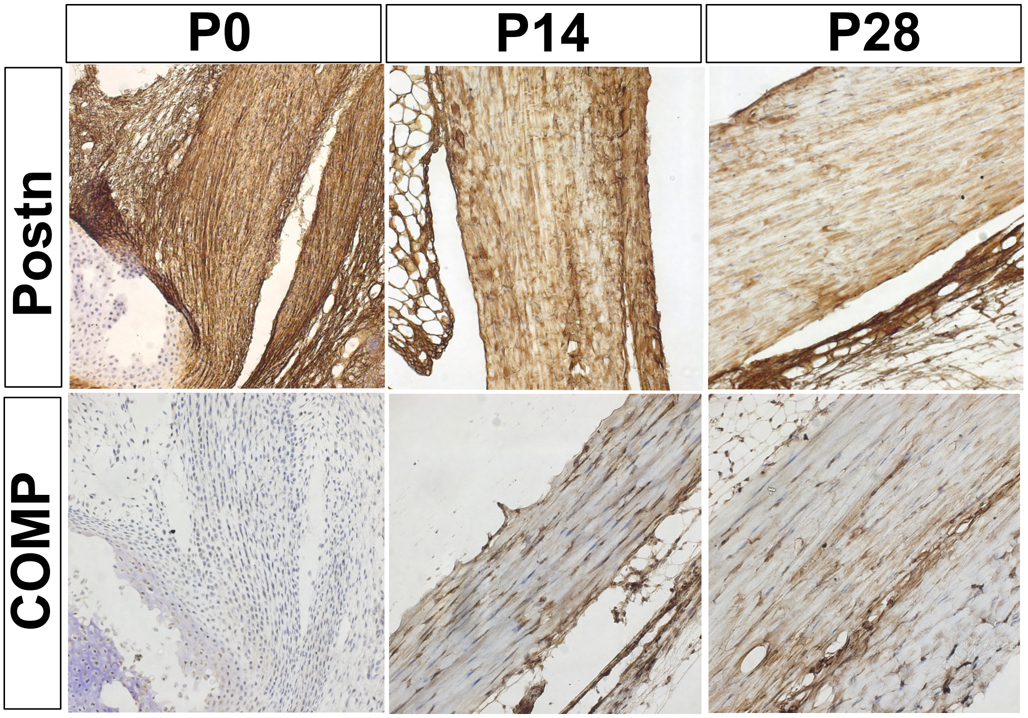

**Supplemental Figure 3**: Immunohistochemistry validation of select proteomic targets Periostin and COMP across developmental timepoints.
